# Revisiting Colocalization from the Perspective of Similarity

**DOI:** 10.1101/2022.05.23.492977

**Authors:** Luciano da Fontoura Costa

## Abstract

Given two or more concentrations, an interesting and important related issue concerns the quantification of how strongly they are spatially interrelated. The concept of colocalization has been frequently considered as an indication of the tendency of the values of two concentrations to spatially vary together. While this frequently adopted approach presents several interesting characteristics, being a suitable choice for several situations, in the present work we study how multiset similarity indices can be applied for similar purposes, possibly allowing a complementation, in the sense of taking into account shared portions of the concentrations, of the colocalization characterization provided by the Pearson correlation methodology. The problem of colocalization is first addressed in terms of possible underlying mathematical models, and then the Pearson correlation coefficient-based approach, as well as the standardization procedure which is its intrinsic part, are presented and discussed. The particularly important issue of how to define the baseline of the concentrations is also approached and illustrated. The minmax alternative normalization scheme is presented next, followed by the description of the three considered multiset simiarlity indices — namely the interiority, Jaccard, and coincidence similarity approaches. The characteristics of each of these methods is then illustrated respectively to 1D, and then to 2D concentrations under presence of several interesting and relevant effects including spatial displacement, as well as sharpening, presence of unrelated effects. The similarity indices, and in particular the coincidence approach, are found to present some interesting features when applied to the quantification of the colocalization between two or more concentrations, suggesting that it can provided complementary information when performing colocalization analysis.

## 1 Introduction

With the advancements in molecular biology experimental resources, in particular fluorescence imaging, it has become possible to mark in accurate and systematic manner gene expression and other biochemical concentrations in tissues and organs. As these advances continue, increasingly more comprehensive and accurate results are obtained, typically in the form of images of densities or concentrations, that demand more and more accurate and effective means for their respective analysis in terms of objective quantifications of the relationship between two or more densities.

The characterization and analysis of relationships between co-existing biochemical densities and concentrations has been approached predominantly through *colocalization analysis*, often being related to the application of the Pearson correlation coefficient between the respective values (e.g. [1, 2, 3, 4, 5]). While these methods are interesting on themselves and are therefore posed to continue to be employed, it becomes interesting to consider alter-native approaches that can complement the characterization of coexisting concentrations and densities. This constitutes the main motivation of the present work, which adopts the concept of multiset (e.g. [6, 7, 8, 9, 10, 11]) similarities — more specifically the interiority (or overlap [12]), Jaccard (e.g. [13, 14, 15, 16, 17]) and coincidence indices (e.g. [18, 19, 20]) — as means for quantifying the colocalization of two or more densities.

In the present work, we will consider *concentrations* as not being necessarily normalized to unit area/volume, white the terms *density* or *distribution* is applied to concentrations which have been otherwise normalized.

We start from the principle that the *colocalization* of two densities means that the respective biological effects tend to take place at the same parts of the considered tissue or organ. This important problem is illustrated in Figure 2, involving two hypothetical densities imaged as (a) and (b). In this case, their combined visualization in (c) readily indicates that, though the two densities are moderately colocalized, they are indeed characterized by a marked spatial shift that was emphasized by the joint visualization.

**Figure 1:**
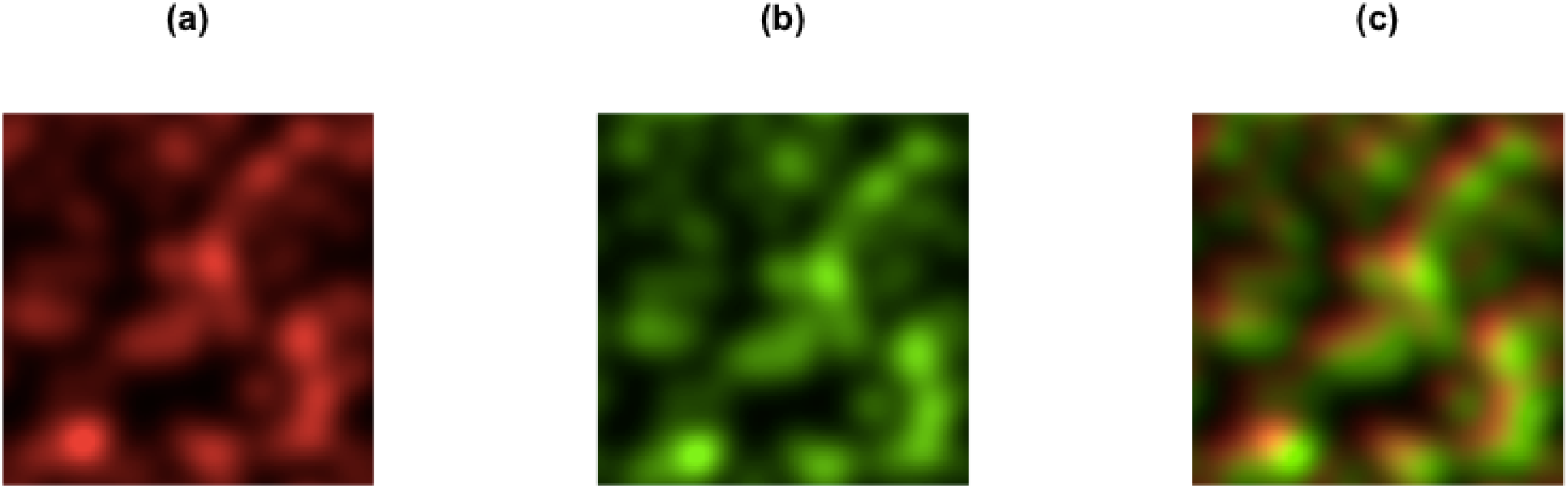
An illustration of the colocalization task. Two distinct biochemical densities are acquired as two respective images (a) and (b). Figure (c) illustrates the image obtained by superimposing the two expressions so that the colocalized portions tend to appear as yellow. In this particular case, it can be readily observed from (c) that the two densities in (a) and (b), though moderately colocalized, also involve a significant relative spatial displacement.

**Figure 2:**
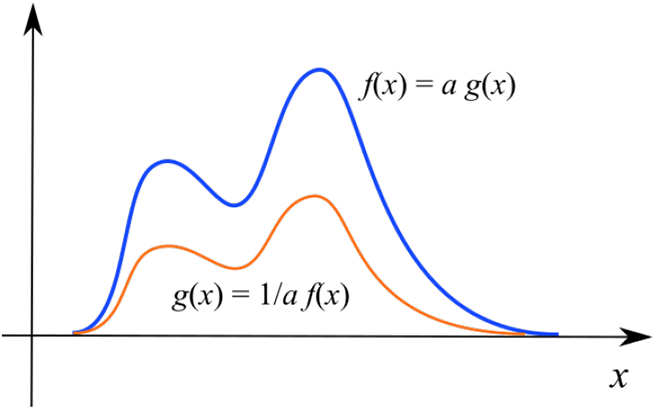
The proportionality relationship that, in the present work, is understood to underly the concept of colocalization (a). Two linear concentrations *f* (*x*) and *g*(*x*) are not equal, but any of them can be obtained by multiplying the other by a specific factor *a*. The different heights indicate that the two concentrations were generated at distinct intensity. Biologically, the two concentrations could correspond to proteins that are being expressed at the same places, but with distinct rates. It may happen that the colocalization is not proportionally observed throughout all the imaged domain, as illustrated in (b), in which there is a region of the domain (larger values of *x*) at which the concentration *f* (*x*) is not being followed by the other concentration *g*(*x*).

In the present work, we employ the *interiority, Jaccard*, and *coincidence* similarity indices in their multiset version (e.g. [18, 19, 20]) as means for quantifying the colocalization between two or more densities. These multiset similarities, in particular the coincidence index, have been found to present special properties that contribute significantly to their selectivity, sensitivity when comparing similar patterns, as well as robustness to localized features perturbations. These properties have paved the way to effective use as basis for several concepts and applications (e.g. [19, 21, 22]).

This work starts by addressing the important issue of expressing colocalization more objectively in terms of mathematical models, with emphasis on proportionality relationships. Then, the traditional colocalization approach employing the Pearson correlation coefficient is presented, illustrated and discussed including the standardization of the concentrations that is always implemented as part of the Pearson-based approach. This is followed by the description of the minmax normalization as well as the three multiset similarity indices considered in this work. Several examples are presented illustrating the respective characteristics of the several approaches here considered for colocalization quantification, first respectively to 1D concentrations, and then taking into account several 2D cases.

## 2 Colocalization Models

Before proceeding to quantifying colocalization, it is important to have an objective comprehension of what it is. Probably the best approach to this problem consists in identifying some *mathematical models* that can be used to define, or at least to characterize, what it will be meant by colocalization. This interesting issued is developed in the present section.

Literally, the term *colocalization* plainly means that something is occupying, or tending to occupy, the same positions than another thing. For instance, in biology, the term is often employed to mean that two protein types are being expressed at the same places. Thus, we should expect that one of the concentrations will tend to follow the other, in the sense that when one is large/small the other will also be or, in other words, that one of the concentrations will be *proportional* but not necessarily identical to the other, which can be mathematically expressed as:

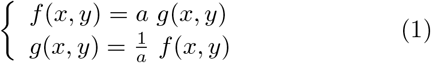

where *a* ∈ ℛ^+^ is a non-negative generic multiplying factor.

It is also possible, for generality’s sake, to incorporate a constant term *b* as:

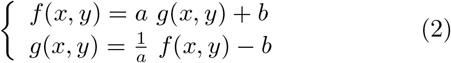

Figure 2 illustrates the concept of colocalization in terms of proportional relationship between two one-dimensional concentrations *f* (*x*) and *g*(*x*).

This relationship between the two concentrations, which we will understand as reflecting this concept in the most literal manner as possible, therefore yields a linear relationship between the two concentrations, as illustrated in Figure 3 respectively to two one-dimensional concentrations *f* (*x*) and *g*(*x*).

**Figure 3:**
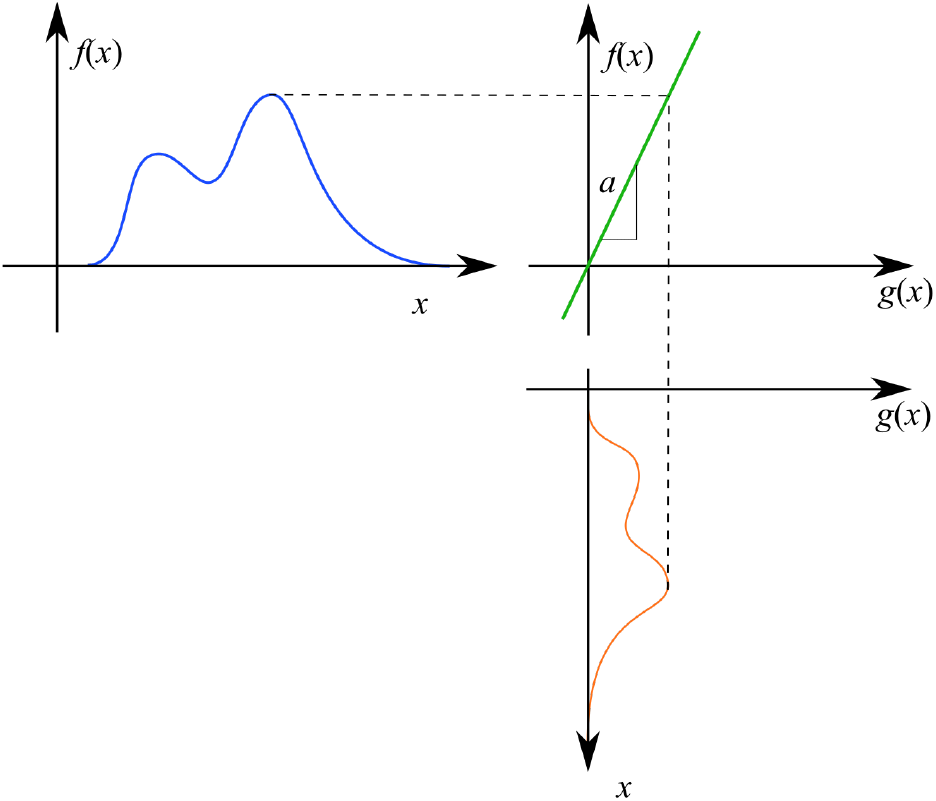
The basic understanding of colocalization adopted in the current work consists in assuming that one of the concentrations, *f* (*x*) is proportionally related to another concentration *g*(*x*) by a proportionality factor *a*, as illustrated in this figure for the case *a >* 1.

It is interesting to observe that the concept of colocalization as formalized above is not necessarily the same as the idea of *identity* between two concentrations, given that two colocalized profiles are not necessarily identical. In case the concept of *similarity* does not allow for concentration scalings, it will also become distinct from the idea of colocalization.

In practice, several effects can eventually influence the basic proportionality assumption in the colocalization. For instance, we can have:

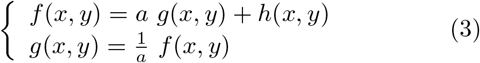

For instance, *h*(*x, y*) can be a uniformly random noise field, or it may be caused by another independent effect not related to *g*(*x, y*). The latter possibility is illustrated in Figure 4. Here, we have the same proportional concentrations already seen in Figure 3, but with the difference that an additional peak appears in the concentration *f* (*x*). This peak is not related to *g*(*x*) in any direct manner, being a consequence of an additional, independent influencing factor.

**Figure 4:**
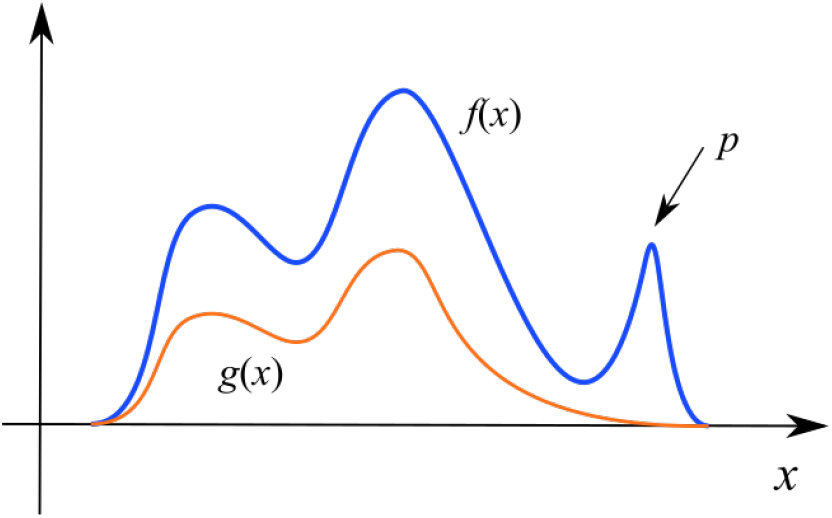
Two concentrations characterized by being generally proportional except for the peak *p* appearing on the right portion of concentration *f* (*x*).

Another interesting situation to be taken into account when analyzing colocalization appears when one of the concentrations corresponds to a shifting of the other, i.e.:

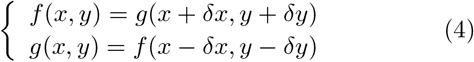

It is also possible that one of the concentrations undergoes modulation by another independent field *h*(*x, y*), such as corresponding to a gradient along the concentration domains, e.g.:

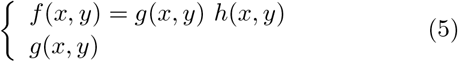

The concentrations can also be related through function composition, i.e.:

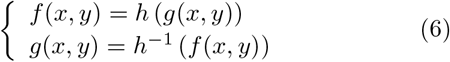

As an example of the above effect, we have the situation in which each of the concentrations correspond to the reciprocal of the other:

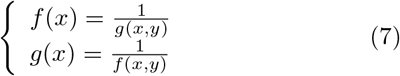

Needless to say, the several mathematical representation schemes above can appeared combined in their several forms. The planning, quantification, and interpretation of colocalization applications can benefit substantially in case the respectively underlying mathematical model is known or hypothesized (and then subjected to validation).

## 3 Standardization and the Pearson Correlation Coefficient

The average value of a 1D continuous concentration *X*(*x*) is defined as:

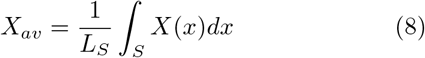

where *S* is the support of the concentration, i.e. the interval of *x, In* = [*x*_*m*_, *x*_*M*_], for which the concentration is of interest for our analysis. The quantity *L*_*S*_ refers to the extension of the interval *In*.

For 2D concentrations, the following average definition applies

In case the concentration is sampled in terms of *N* points, sampled at equally spaced intervals, we have that its average value concentration *X*_*i*_, *i* = 1, 2, …, *N*, can be estimated as:

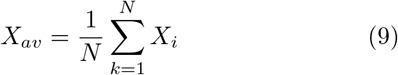

In the case of a concentration expressed as:

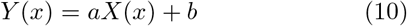

its average can be found to be equal to:

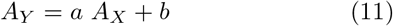

Given *N* samples *X*_*i*_, *i* = 1, 2, …, *N*, of a random variable *X*, they can be *standardized* (e.g. [23, 24, 25]) by making:

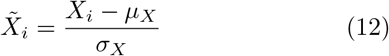

where *µ*_*X*_ and *σ*_*X*_ are respectively the mean and standard deviation of *X*. The standardized variables 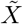 have some special properties, including null means and unit standard deviation. In addition, most of them result comprised in the interval [−2, 2]. Though these properties are often useful and adopted, it should be borne in mind that information about the original variable is lost in the process, especially concerning its original average value. We will come back to this respectively to the Pearson correlation coefficient.

Given two sets of observations *X* and *Y* of two respective random variables, their *Pearson correlation coefficient* (e.g. [26, 27, 28, 23, 25])) can be estimated by performing the two following steps: (1) standardize each set separately; and (2) calculate the following average:

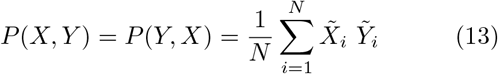

It can be verified that −1 ≤ *P* (*X, Y*) ≤ 1.

The Pearson correlation coefficient quantifies the tendency of the two variables *X* and *Y* to vary together. For instance, when one increases, the other also tends to increase. Similarly, when one decreases, the other tends to accompany the trend. The higher the value of the Pearson correlation coefficient, the strongest this tendency is. Negative values of Pearson correlation coefficient indicate that when one of the variables increases, the other decreases, and vice-versa.

Given the above characterized operation of the Pearson correlation coefficient, it is intrinsically suitable for expressing the colocalization between two concentrations *X* and *Y*, being therefore amply adopted. An important aspect of the Pearson correlation coefficient to be considered at all times concerns the fact that its calculation involves the standardization of the two variables, therefore implying the loss of some original information including the averages *µ*_*X*_ and *µ*_*Y*_ of the two variables. For instance, consider the particularly interesting situation depicted in Figure 5.

**Figure 5:**
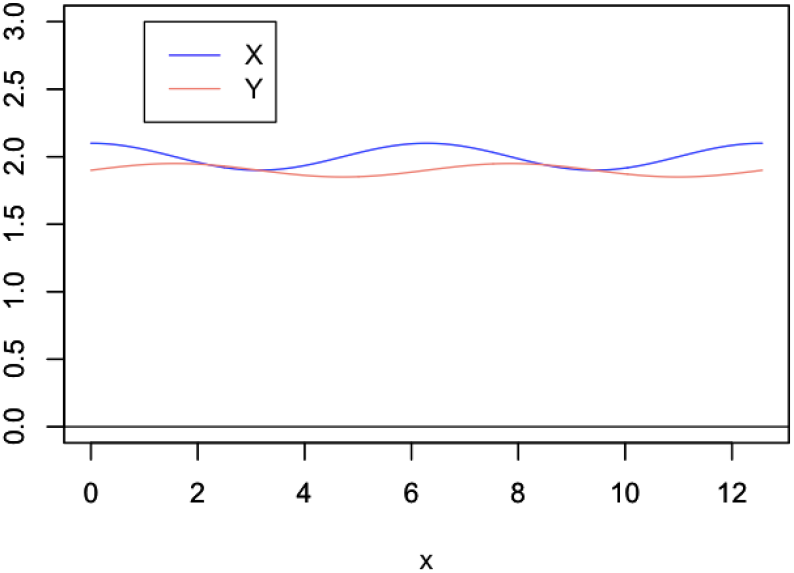
Two one-dimensional concentrations *X*(*x*) and *Y* (*y*) have average values approximately equal to 2.0, but both these concentrations incorporate some small scale waving. Despite the overall similarity between these two concentrations, their respective Pearson correlation coefficient can be verified to be zero.

The Pearson correlation coefficient for these two concentrations is zero because the original averages are not taken into account and also because the respective small scale oscillations correspond to a sine and a cosine. If, instead, we had to sines or cosines of any positive amplitude, the Pearson correlation coefficient would be equal to 1. Thus, we have that small changes in small scale detail of the concentrations can make the Pearson correlation to change from 0 to 1 (or even -1). In case we are interested not only on the joint variation between the concentrations, the original averages can be estimated before standardization in order to be considered later.

Let us now present another example of quantifying the colocalization of the two signals in Figure 6. These two concentrations were defined from a reference concentration *Z* as:

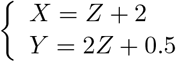

The standardization of the concentrations will remove the respective averages and rescale the results so that they have the same dispersion. These combined processing effectively yields 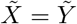, therefore implying the respective Pearson correlation coefficient to result equal to its maximum value of 1. This is completely reasonable because the two concentrations were built so as to vary together proportionally.

**Figure 6:**
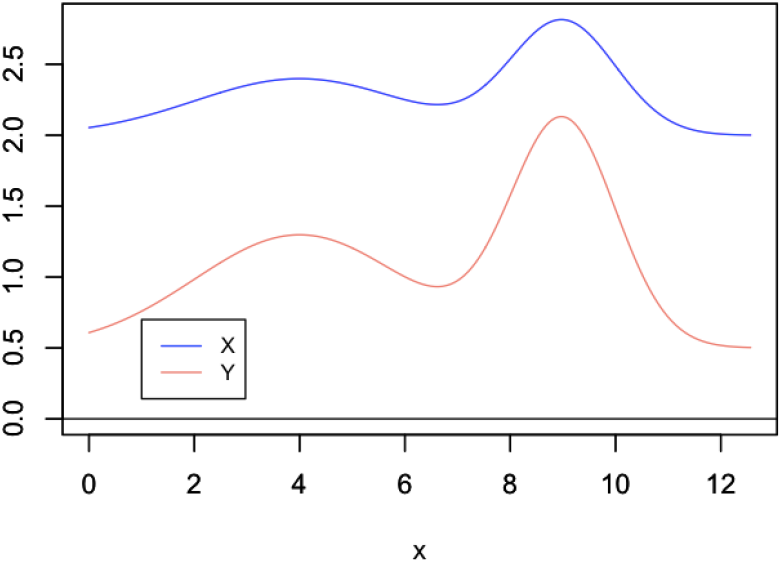
Two concentrations that have markedly distinct averages and magnitudes, but which have shapes proportional one to the other. As a consequence of the standardization implemented in the Pearson correlation coefficient, its value results 1 for this specific example.

There is another type of concentration representation that deserves special attention in colocalization studies, namely the situation in which the concentrations have area (in the case of 1D) or volume (2D) equal to one, in which case they can be understood as *probability densities*. Given any non-null concentration, it can be transformed into a respective probability density function simply by dividing each of its values by the total area *A* or volume *V* (assuming unit sampling intervals). The effect of this transformation on the proportional relationship here adopted as a reference for understanding colocalization then becomes:

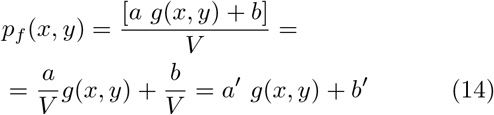

Therefore, the translation of a generic concentration into its respective probability density form does not affect, other than proportional changes on the involved coefficients, the proportionality relationship.

Given the importance of the effect of the average level of the concentrations on the localization, we now discuss another related example illustrated in Figure 7(a). We have both concentrations characterized by a smaller waving pattern on a larger oscillation. More specifically, the concentrations have been defined as:

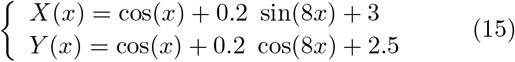

The scatterplot and respectively obtained Pearson correlation coefficient are shown in Figure 7(b).

**Figure 7:**
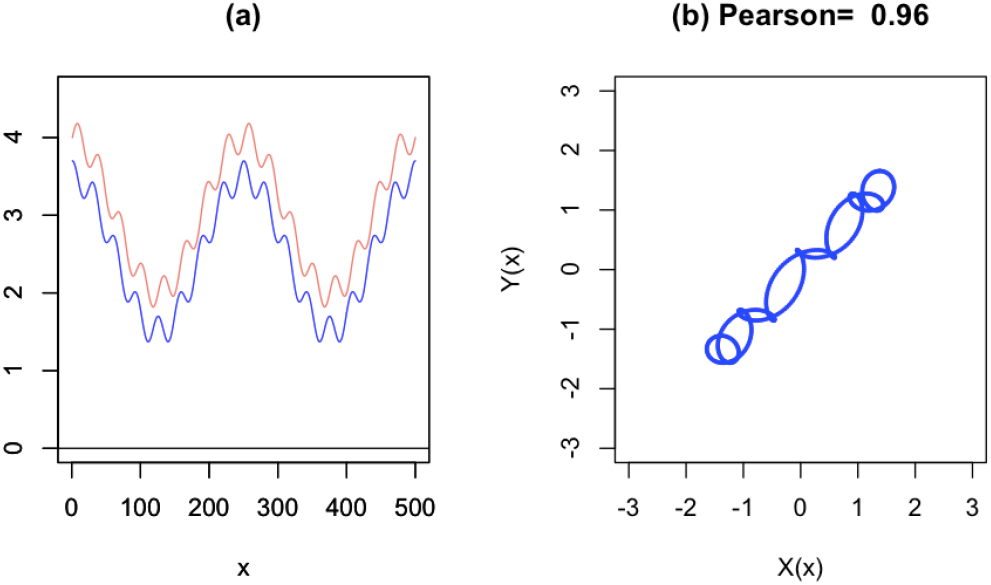
Two concentrations presenting two scales of oscillations (a) and the respective scatterplot and Pearson correlation coefficient (b), which can be understood to characterize the colocalization at the largest spatial scale of the two cosines cos(*x*). The null colocalization existing between the two smaller oscillations cannot be characterized directly by the Pearson correlation coefficient, demanding the prior subtraction of the larger cosine baseline.

The relatively high value obtained for the Pearson correlation coefficient can be understood to quantify the colocalization of the larger oscillation parts of the two concentrations. However, in case these larger oscillations are understood as a baseline, with the smaller (in terms of height) oscillations patterns, namely sin(8*x*) and cos(8*x*), therefore constituting the pattern of interest, a null Pearson correlation value should have been obtained. In the case of the Pearson correlation, this could be achieved by subtracting the larger oscillation terms from both concentrations prior to respective calculation. Though in most practical cases the functional description of the concentrations are not available, there are mathematic-computational methods that can be used for estimating the baselines of the two concentrations as corresponding to the larger oscillation.

In this work, the baseline is understood to mark the region of interest in the concentrations. More specifically, the portions of the concentrations that are above or around the baseline are to receive special attention in the analysis. For instance, if interest is focused on the smaller variations taking place on larger oscillations or even a constant plateaux, the baseline can be set so as to keep those portions apart from the analysis.

Figure 8 presents the same concentration *X*(*x*) as above as well as five possible choices, amongst the infinite possibilities, of baselines. The black baseline coincides with the *x*-axis, therefore providing a natural reference in case the null concentration is to be taken as reference. The concentration in brown, corresponds to the average of the concentration. The green baseline is defined by the minimum of the concentration. The magenta baseline is an arbitrary choice possibly motivated by specific interests of the research. While all these three baselines correspond to constant values, the baseline in red is a cosine function that has been defined from the larger oscillation in the concentration.

**Figure 8:**
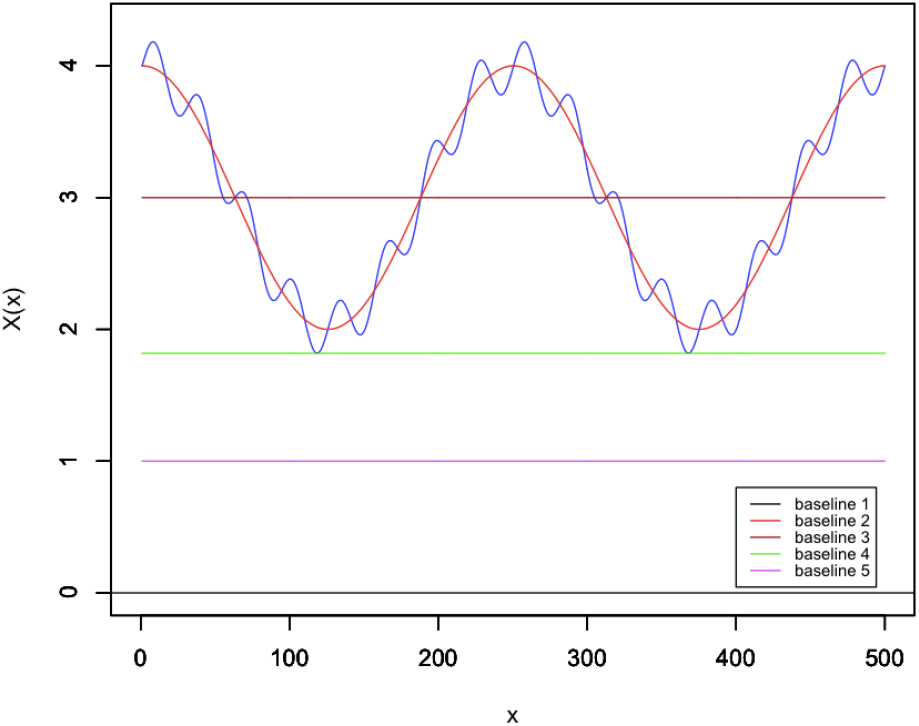
A given concentration *X*(*x*) (in blue) and five possible choices of baselines. Only the brown baseline is effectively considered by the Pearson correlation approach. The red baseline can also be considered for the Pearson analysis provided it is subtracted beforehand from the concentration.

Importantly, the Pearson correlation approach can only take into account the brown baseline, as it corresponds to the average of this signal and the standardization shifts the concentration always to this position. The oscillating baseline can also be considered in the Pearson analysis provided it is subtracted from the concentration first. The other constant baselines cannot influence the Pearson correlation coefficient in any way as the standardization always place the concentrations at the respective average. An important conclusion therefore is that the result of Pearson correlation analysis frequently depends substantially on the definition of the baseline. Two more evident possible baselines in the case of the previous example correspond to: (i) the overall average of the concentrations; and (ii) the larger oscillation itself. However, there is an infinite additional number of choices for baselines, each to be chosen among the infinite possible functions defined in the same interval of *x* of interest. Importantly, the Pearson correlation cannot cope with constant baselines along that interval, because this term would be otherwise removed by the standardization that is intrinsic to any Pearson correlation coefficient calculation.

We can conclude from the above reasonings that the standardization-Pearson analysis will be particularly suitable for the analysis of concentrations that can be expressed in the following general form:

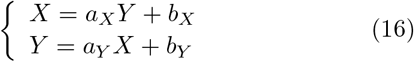

except for any choice of constant baseline.

Some other types of relationships can be eventually transformed into the above representation through mathematical manipulations involving taking logarithms, etc. In addition, the above relationship can be readily generalized to more than two concentrations, e.g.:

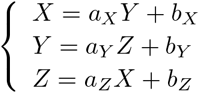

Henceforth, the constant terms in these expressions (e.g. *b*_*X*_, and *b*_*Y*_ in Eq. 16), which do not necessarily correspond to the average value of the respective concentrations (as this can also be influenced by the other terms), will be henceforth referred to as *constant terms*.

Let us now consider a colocalization case, depicted in Figure 9, in which one of the concentrations has a well-defined peak that is not related in any means to the other concentration. More specifically, in this case both concentrations also have the same area, so that we would have half of the area of the concentration *X* colocalizing with the other concentration. However, the obtained Pearson correlation coefficient resulted in *P* (*X, Y*) = 0.8007, which suggests a colocalization much higher than the half of shared areas.

**Figure 9:**
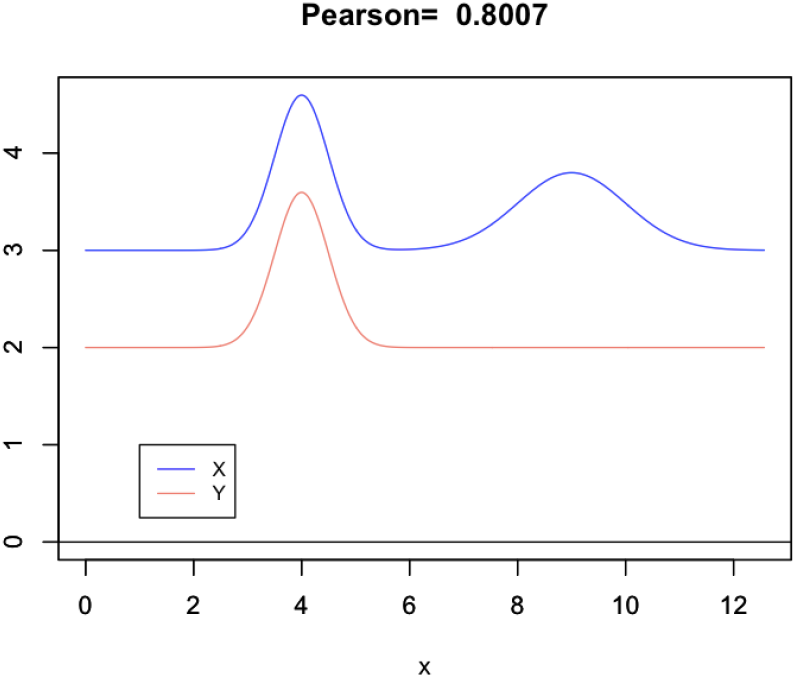
The Pearson correlation coefficient applied to the quantification of the colocalization between these two concentrations, one of which incorporating a peak unrelated to the other, yields a substantially high value. Indeed, one half of the area of the concentration *X* undergoes joint variation with *Y*.

The reason for this surprisingly high Pearson correlation coefficient can be better understood by referring to Figure 10, which shows the standardized versions of the two concentrations as well as the respectively defined scatterplot. It can be verified from the latter that the peak of *X* that is unrelated to *Y* yields a much shorter spoke in the scatterplot that, in addition to the displacement of the scatterplot pattern respectively to its coordinate origin, results not being enough to reduce more substantially the otherwise estimated Pearson correlation coefficient.

**Figure 10:**
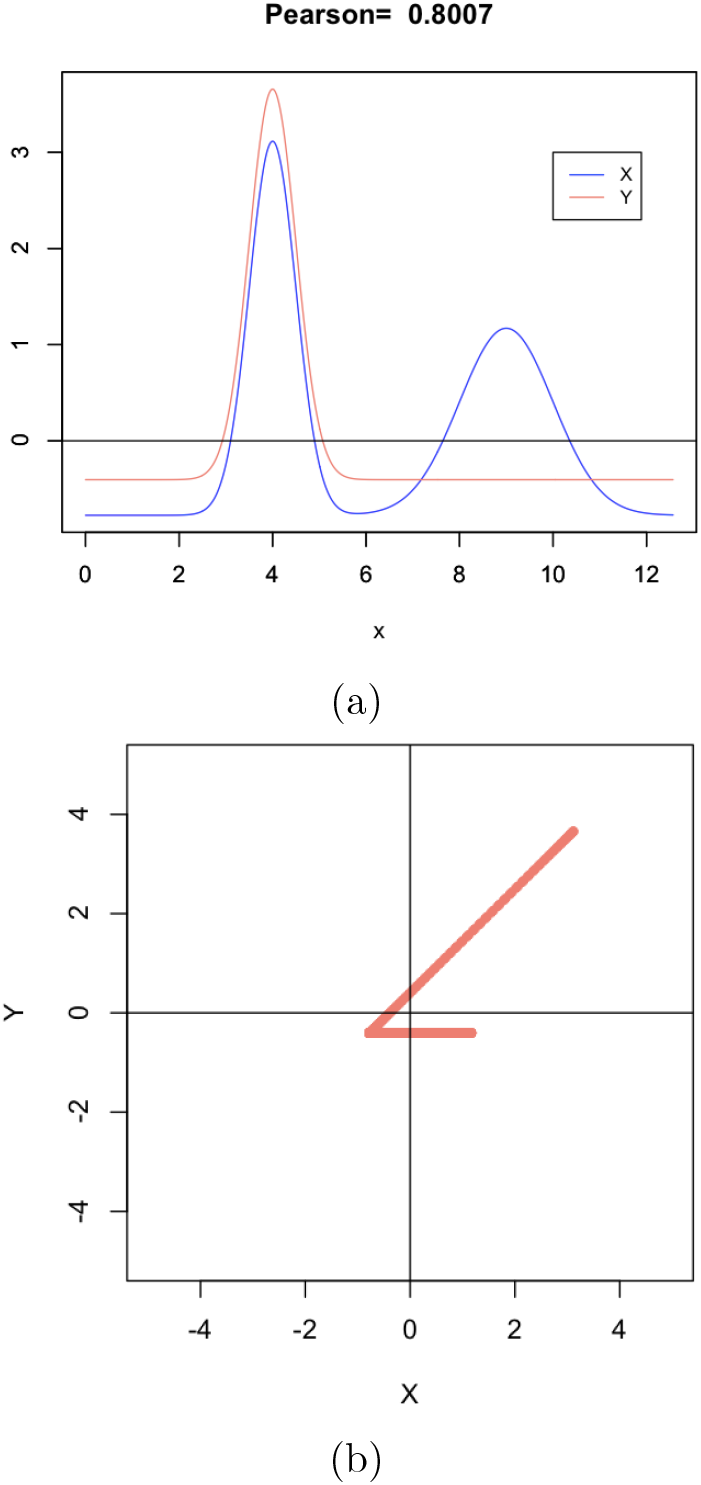
The standardized versions of the two concentrations in Figure 9(a), as well as the respectively obtained scatterplot, which reveals the predominance of the joint variation of just one of the peaks of the concentration *X* with the colocalized peak along *Y*. The additional, unrelated peak in *X* defines a much shorter spoke in the scatterplot that is not sufficient to reduce the Pearson correlation to a smaller value.

All in all, though the standardization of the original concentrations provides a pre-processing that tends to be suitable and compatible with subsequent colocalization analysis by employing the Pearson correlation coefficient, there are situations such as that illustrated above that can be understood as an overestimation of the colocalization. In the following section we consider potential of three multiset similarity indices for possible addressing the overestimation issue.

## 4 Minmax Normalization and Multiset Similarity Indices

Another interesting approach to normalizing the two concentrations to be analyzed is by using a method that in this work will be referred as *minmax normalization*. This can be readily implemented as:

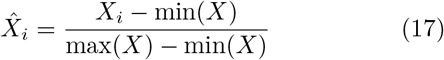

As a consequence of this normalization, the values of *X* become all comprised between 0 and 1. When preparing the concentrations for subsequent colocalization analysis, the minmax is applied independently on each of the involved concentrations.

From the biological point of view, this normalization can be understood as removing the constant expression term and then normalizing the maximum expression to 1. Also known as *overlap* (e.g. [12]), the *interiority* index between two concentrations quantifies how much one of the concentrations is contained into the other. It can be expressed as:

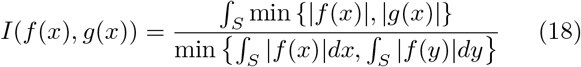

with 0 ≤ *I*(*f* (*x*), *g*(*x*)) ≤ 1 and *I*(*f* (*x*), *g*(*x*)) = *I*(*g*(*x*), *f* (*x*)).

The real-valued Jaccard similarity index between two concentrations can be written (e.g. [18, 19]) as:

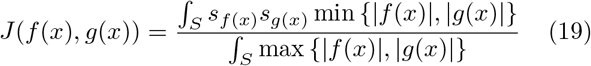

where *s*_*f*(*x*)_ = *sign*(*f* (*x*)) and *s*_*g*(*x*)_ = *sign*(*g*(*x*)). We also have that −1 ≤ *J* (*f* (*x*), *g*(*x*)) ≤ 1.

The Jaccard index quantifies how similar the two concentrations are while considering their shared area and outline [20].

Given that the Jaccard index cannot account for the relative interiority between the two concentrations, the *coincidence similarity index* has been proposed [18, 19] as the product of the Jaccard and interiority index, i.e.:

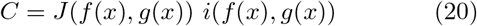

with −1 ≤ *J* (*f* (*x*), *g*(*x*)) ≤ 1 and *C* = *J* (*f* (*x*), *g*(*x*)) = *J* (*g*(*x*), *f* (*x*)).

These indices, and in particular the real-valued Jaccard and coincidence cases, present some intrinsic interesting features [18, 19, 20] that make them particularly selective, sensitive and robust to localized features perturbations. These characteristics have allowed successful applications of these indices to several areas and types of problems, including template matching [19, 22] and translating datasets into networks [21].

In the present work, we study how these three similarity indices can be applied as an alternative methodology for characterizing the colocalization between two given concentrations. Interestingly, the three similarity indices above can be applied with or without preliminary standardization of the concentrations, depending on the application requirements and type of data to be analyzed. The ability of these indices to cope with constant baselines other than the concentration average constitute on aspect of particular interest for colocalization analysis.

In the case standardization, the obtained similarity indices can be understood, in analogous manner to the Pearson correlation characterization, as quantifying the tendency of joint variation presented by the two concentrations. Another interesting possibility consists of applying the minmax normalization, resulting in two non-negative normalized concentrations with non-null average values, in which case the similarity values will relatively indicate the shared area (for 1D concentrations) or volume (for 2D concentrations) between the two concentrations. A third possibility consists of implementing no normalization, so that generic constant baselines can be specified.

Some of the possible pre-processing that can be applied to the concentrations before similarity colocalization are summarized in the following. Only alternatives B and C apply to the Pearson correlation colocalization, as it always implements standardization.

### A No standardization

The concentrations are used as given;

### B Baseline subtraction

Subtract a baseline of particular interest from the concentrations. This will only have an effect on Pearson correlation-based colocalization in case the baseline does not have a constant value;

### C Standardization

The concentrations are standardized prior to colocalization quantification so as to have average equal to zero and standard deviation equal to one. Most of their values will result within the interval [−2, 2];

### D Area/Volume normalization

Divide each of the concentrations by their respective area/volume so that they will result with unit area (1D) or volume (2D);

### E Minmax normalization

The concentrations are minmax-normalized. Both concentrations will be normalized between zero and one, therefore having the baseline not corresponding to their respective averages, but to the constant zero. Observe that this normalization does not generally yield the same results as the same area/volume normalization.

Recall that, in addition to these possibilities, any baseline of interest can be subtracted from the concentrations as a preliminary step.

Figure 11 illustrates the application of the coincidence similarity for the characterization of the two concentrations in Figure 7(a), both of which incorporating respective non-null constant terms. The results in Figure 11(b) refer to the presentation of the two concentrations by ordering the involved pairwise minimum values and averaging the respective maximum around a window centered at each instance. The Jaccard can then be readily approximated visually as corresponding to the quotient between the area below the minimum curve and the area of the average maximum curve (exact Jaccard estimation is obtained by considering the original maximum values, though they tend to be much more jagged). The type of concentration re-ordering shown in (c) is henceforth referred to as *sorted representation*. The obtained results cannot be directly compared to the respectively obtained Pearson correlation coefficient of *P* = 0.96, since here the concentrations have not been previously standardized, therefore presenting pronounced constant terms that are duly taken into account by all the considered similarity indices.

**Figure 11:**
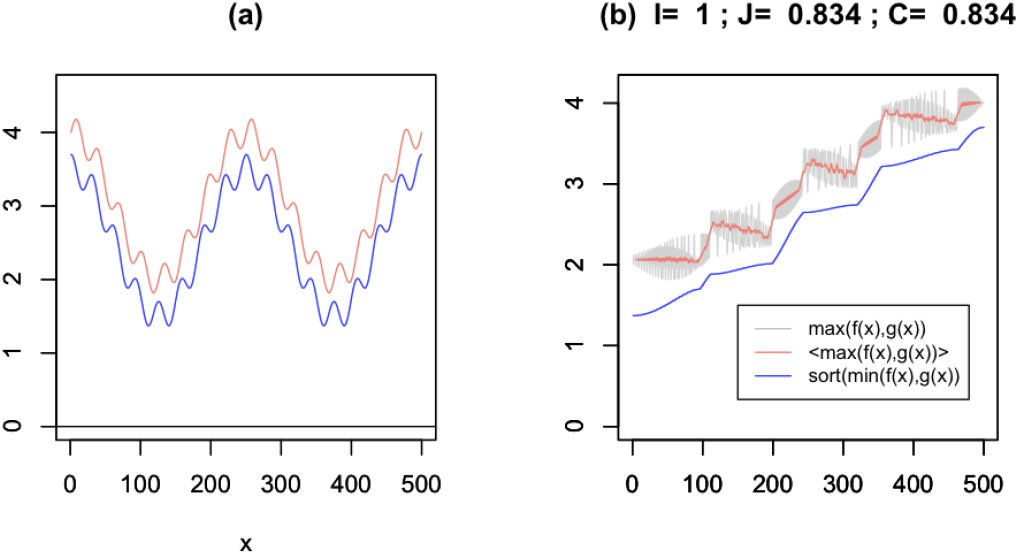
Quantification of colocalization between two concentrations *X* and *Y* involving respective non-null constant terms (a). The respective Pearson is *P* = 0.96. The results respective to the two other indices, namely interiority and Jaccard, are also shown in (b).

Figure 12 illustrates the results obtained by the similarity indices respectively to the same concentrations as in Figure 7. However, now both concentrations were preliminary standardized before the estimation of the similarities. All the three similarity indices yielded colocalization indications smaller than the respective Pearson correlation coefficient *P* = 0.96, which can be understood as reasonable given that the smaller scale oscillations (a sine and a cosine) tend to reduce the colocalization.

**Figure 12:**
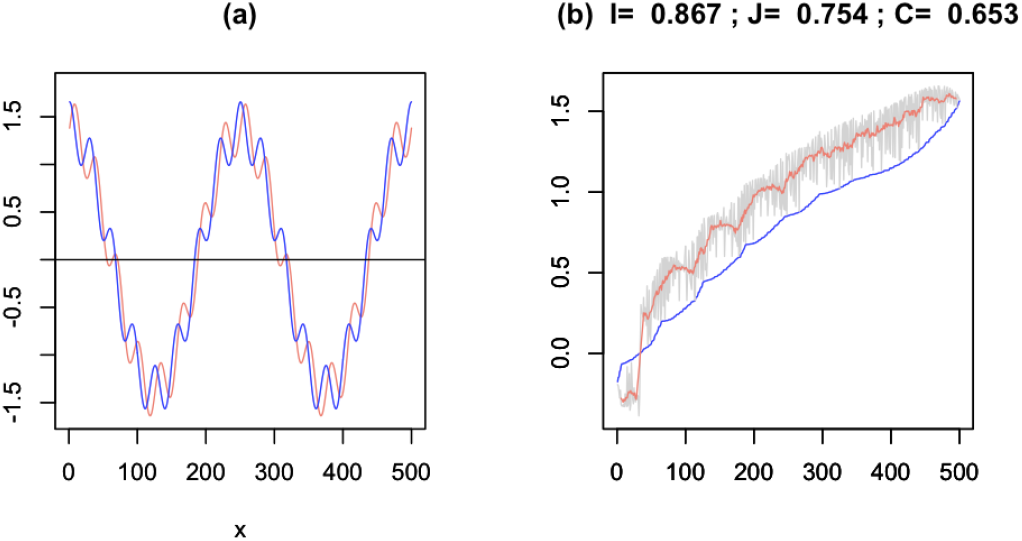
Similarity-based colocalization analysis of the concentrations in Figure 7. Both concentrations were preliminary standardized. All these three indices led to colocalization values smaller than the previously obtained *P* = 0.96, which seem to have overestimated the colocalization.

Figure 13 presents the same situation as before, but now the two concentrations underwent minmax normalization. As expected, the implied higher baseline implied the similarity indices to take into account the shared areas as well as the joint variation tendencies, resulting in moderately larger colocalization values.

**Figure 13:**
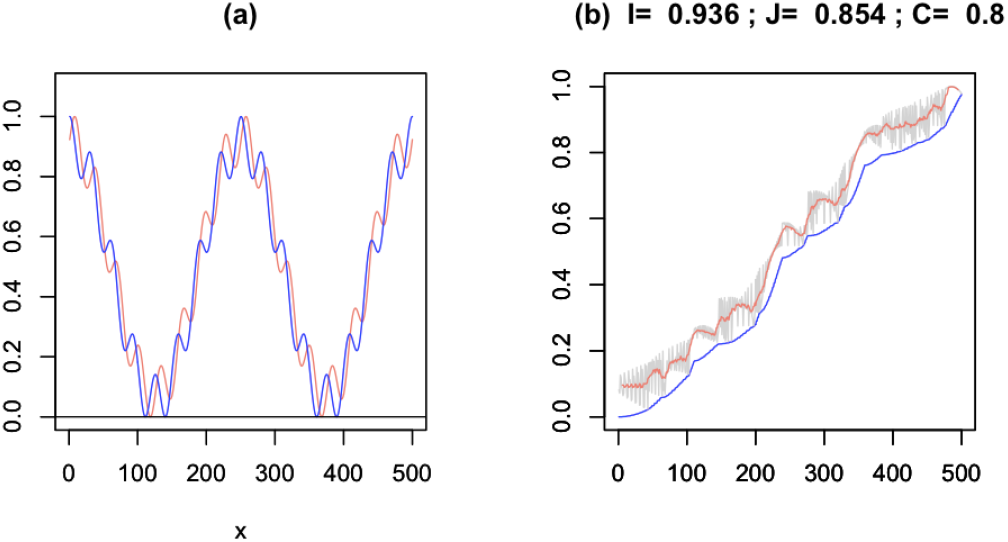
Similarity analysis of colocalization of the the concentrations in Figure 7 adopting minmax normalization.

Figure 14 presents a situation similar to that in Figure 7, but now the two larger oscillation components to both concentrations consist of a sine and a cosine, which do not vary together, leading to low colocalizatiion values provided the concentrations have been previously standardized, as it is the case here.

**Figure 14:**
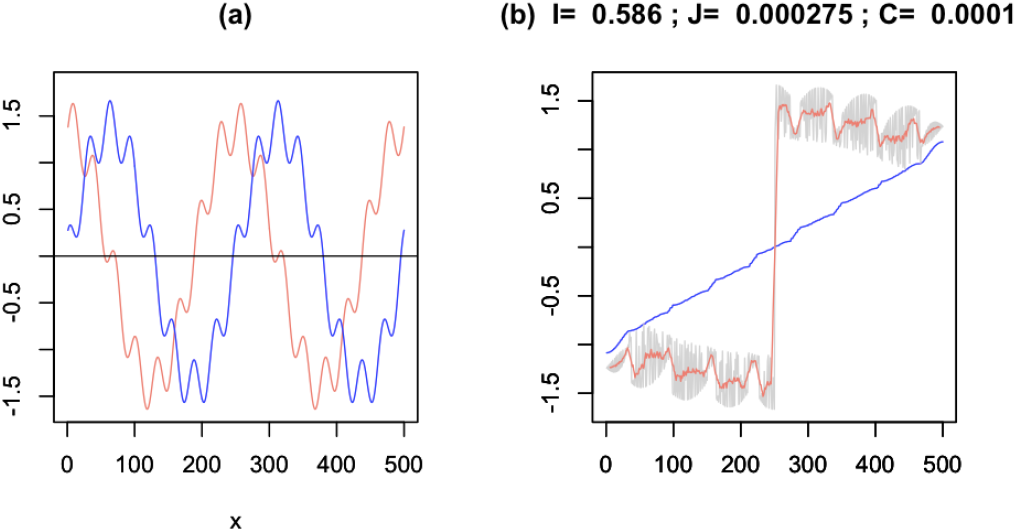
Similarity analysis of colocalization of the two concentrations presenting little net joint variations, obtained while adopting standardization. As could be expected, very small colocalization values has been obtained.

It is important to recall that none of the presented approaches to the analysis of the concentrations in Figure 7 are absolutely right or more suitable. Indeed, the choice of colocalization normalization and index needs to take into account specific specific research and data demands and constraints.

## 5 Colocalization of Noisy in 1D Concentrations

As an illustration of the effect of uniformly distributed noise in the concentrations, we consider the situation in Figure 15, which was obtained from the concentrations in Figure 14 by adding relatively strong uniform noise.

**Figure 15:**
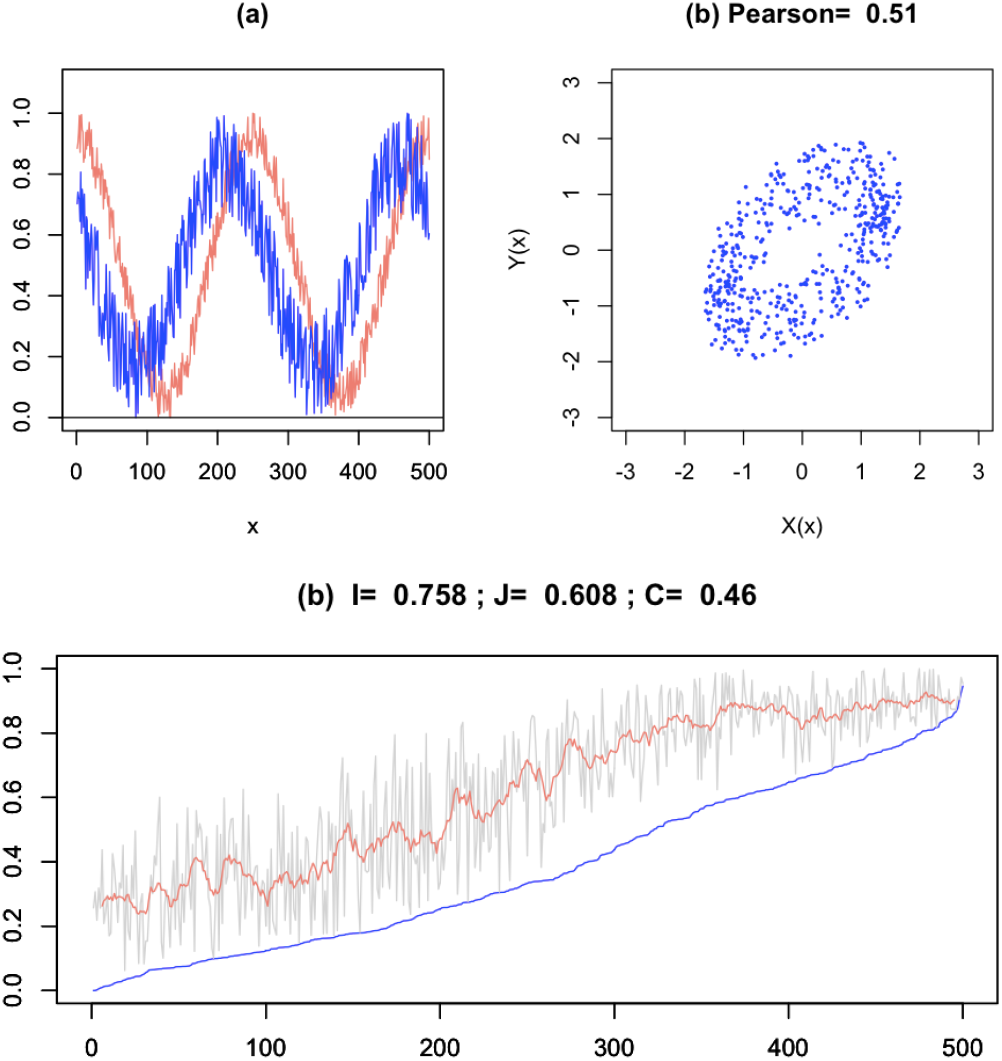
The colocalization analysis of the concentrations in Figure 14 with relatively intense added noise. The colocalization values obtained from all the considered aproaches were substantially reduced.

In this example, the concentrations were preliminary minmax normalized. As it could have been expected, the noisy concentrations yielded smaller colocalization values as gauged not only by the Pearson correlation coefficient, but also by the three similarity indices. By being uncorrelated, uniform noise incorporates similar amounts of joint and opposite variations which tend to cancel one another.

## 6 Colocalization of Hetetogeneous 1D Concentrations

As a example of colocalization quantification in presence of independent expression, we now return to the situation addressed in Figure 10. We had seen that the application of the Pearson correlation coefficient to these concentrations led to a substantial overestimation of the colocalization as a consequence of the additional to one of the concentrations of a peak that had no relationship with the other concentration.

Figure 16 depicts the results obtained by using the multiset similarity index with both concentrations being first minmax normalized. Interestingly, the colocalization value estimated by the coincidence similarity yielded exactly the ratio of the areas of the concentration profile with two peaks with the other concentration. That is a consequence of the fact that the coincidence similarity is based on the shared area between the concentrations, and not only their tendency to vary together.

**Figure 16:**
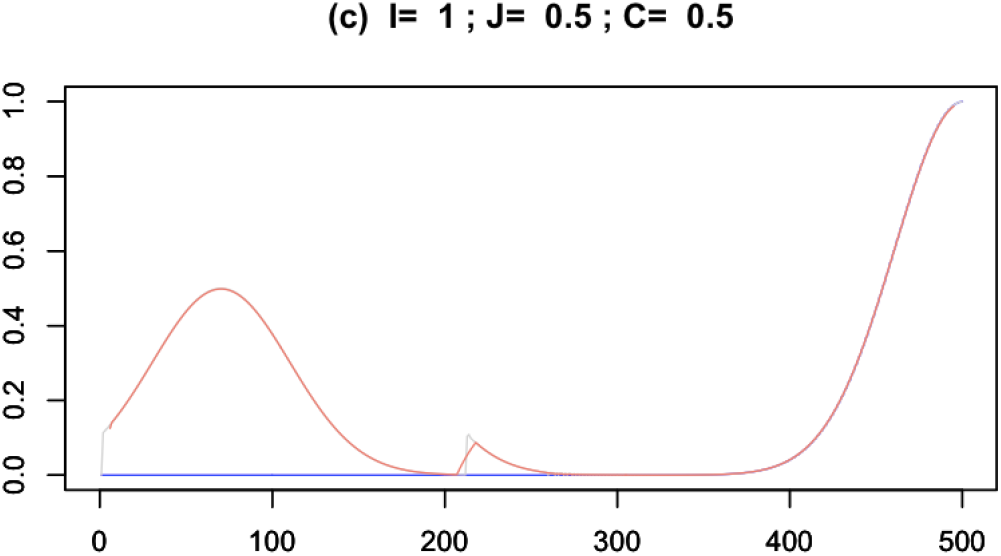
The sorted representation respective to the concentrations in Fig. 10 as employed in the similarity coincidences, which led to more reasonable indications of colocalization in this specific case.

## 7 Colocalization of 2D Concentrations

Having presented and illustrated the possibility to apply the three multiset similarity indices to gauge the colocalization of 1D concentrations, in the following sections we extend these possibilities to several situations involving 2D concentrations. All subsequent results refer to the two concentrations neither having being preliminary normalized before application of the multiset similarity indices. In order to allow complete control on the properties of the concentrations, and also to provide a reference, all 2D concentrations used in this section were synthesized through mathematic-computational methods. More specifically, they correspond to uniformly random spatial distributions of points with given densities that are subsequently smoothed via a gaussian kernel with specific dispersion. All concentrations are then normalized to volume 1, and thus can be understood as probability densities. Figure 17 presents two distinct 2D concentrations that, though being randomly synthesized, still present some considerable overlap, so that some moderate colocalization could be expected. However, the Pearson correlation coefficient yielded the markedly small value of *P* = 0.101. Larger values were obtained from the Jaccard and coincidence similarity values, but with the latter, as could be expected, resulting in the smallest colocalization value between them.

**Figure 17:**
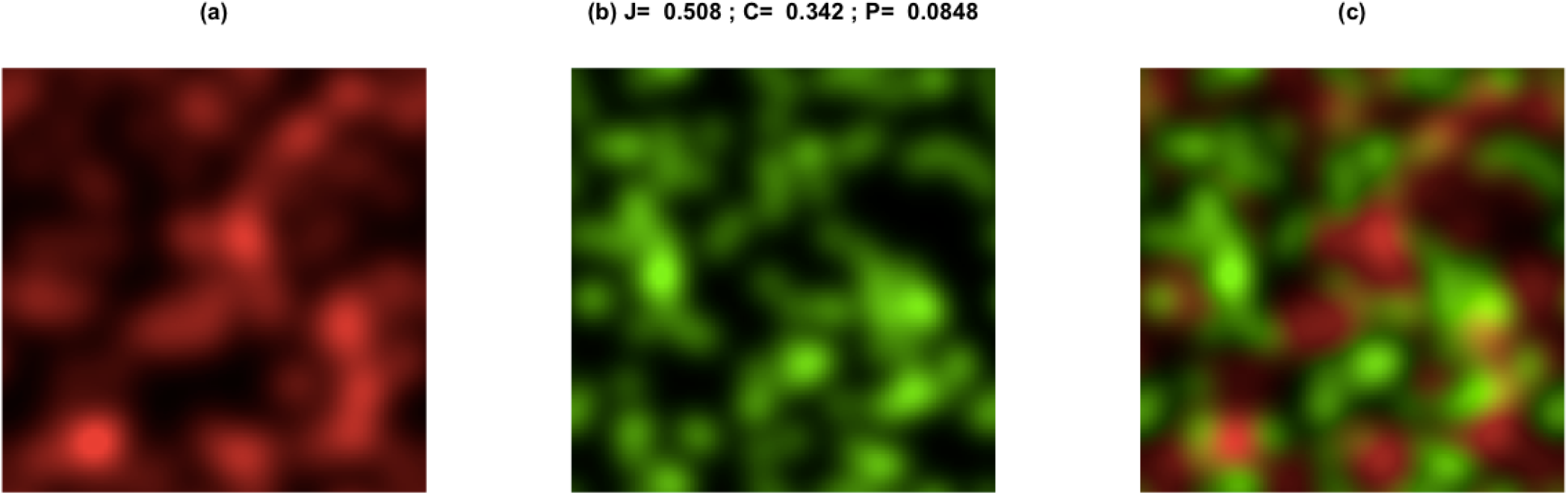
Colocalization of two 2D concentrations randomly synthesized. Though moderate overlap can be observed between these concentrations, the Pearson correlation coefficient resulted markedly small. More reasonable quantifications seem to have been provided by the Jaccard and coincidence similarity indices, with the latter yielding the smallest colocalization value between them.

An interesting situation involving relative spatial displacement, or shift, between the two concentrations is depicted in Figure 18. The two original concentrations were identical prior respectively imposed displacement.

**Figure 18:**
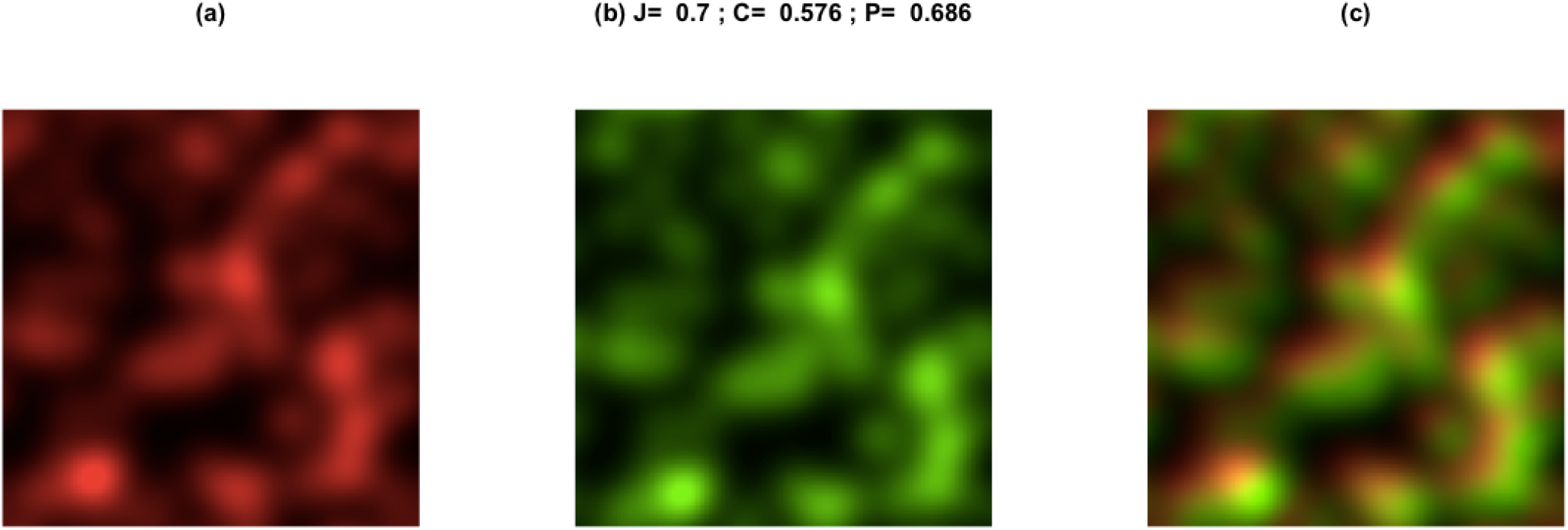
Two identical, but relatively displaced, concentrations. Though a relatively strong displacement can be observed, a Pearson correlation coefficient *P* = 0.686 was respectively obtained. The coincidence-based approach led to a smaller colocalization value.

A particularly interesting colocalization situation arises when one of the concentrations corresponds to a sharpening of the other, as illustrated in Figure 19. The green concentration is so much sharper than the red that it completely predominates in the joint visualization shown in (c), with a good deal of the red concentration not being strongly related to the green. Still, a exceedingly high correlation coefficient of *P* = 0.933 has been obtained. A substantially smaller coincidence value was obtained which seems to better express the colocalization relationship between the two concentrations in this situation.

**Figure 19.**
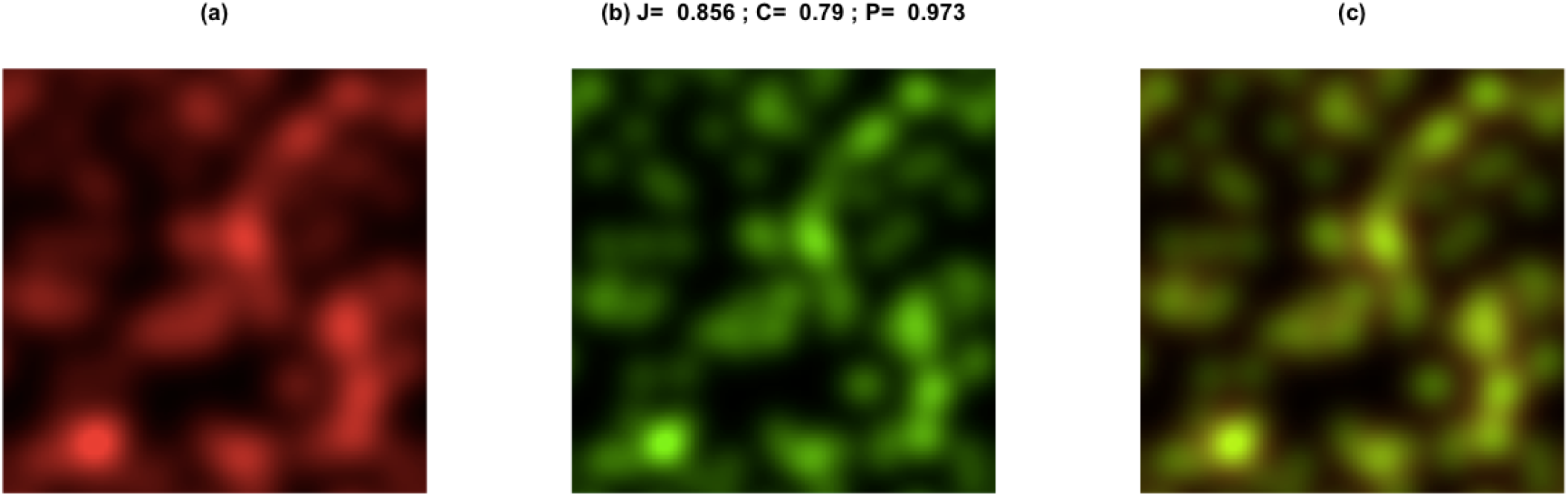

This interesting result stems from the interesting characteristics of the similarity indices in being more accurate when comparing relatively similar patterns [20].

The important situation in which one of the concentration profiles incorporates a markedly distinct part that does not relate whatsoever with the other concentration, as illustrated in Figure 20. Other than the additional spot in the red concentration in (a), both concentrations are identical. Thus, a substantially high colocalization value could be expected, which was the case for the Jaccard and coincidence similarity indices. However, at the same time the Pearson correlation coefficient yielded the result *P* = 0.734, which seems to be too small given that most of the concentrations are nearly perfectly identical other than in the additional spot region.

**Figure 20:**
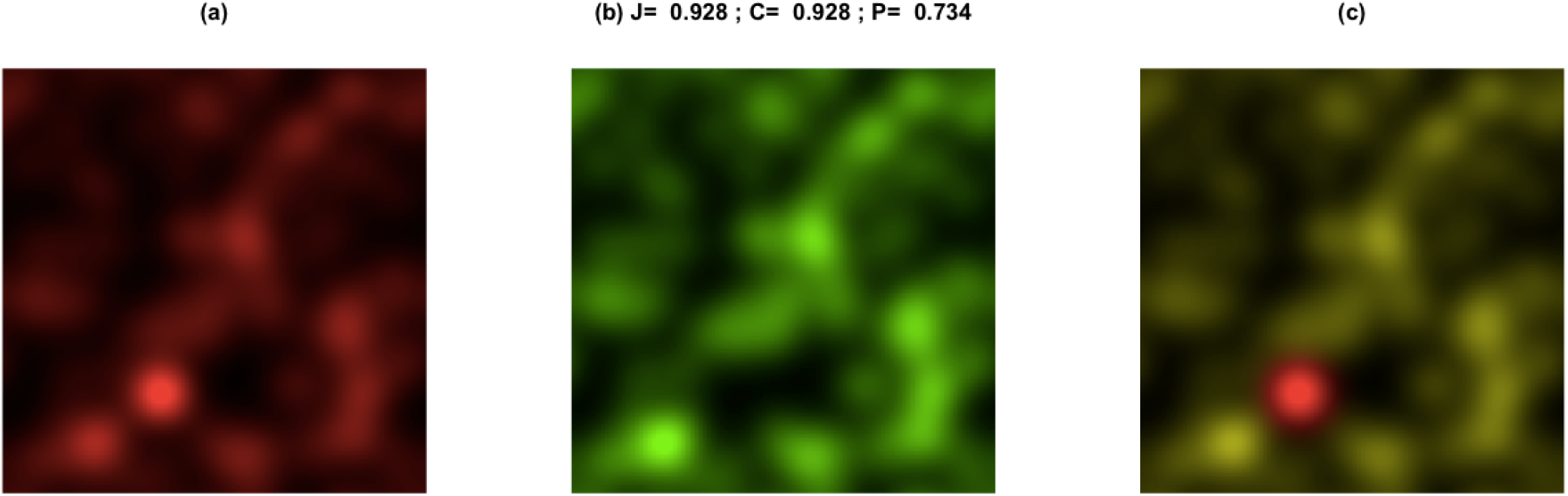
Two identical concentrations, except for the additional spot in the red concentration, which bears no relationship whatsoever with the other concentration. The presence of this localized discrepancy strongly influenced the obtained Pearson correlation coefficient of *P* = 0.734, which resulted markedly small despite the fact that all other portions of the concentrations are in nearly perfect match one another. A more consistent quantification of the colocalization in this particular case seems to have been provided by the Jaccard and coincidence similarity indices.

Figure 21 (a–c)illustrates the colocalization analysis respectively to a situation where one of the concentrations is the reciprocal of the other (Eq. 7). In their own ways, all considered colocalization approaches indicate that the two concentrations are not related: while a negative Pearson correlation was obtained, the Jaccard and coincidence indices resulted very nearly zero. That is mostly a consequence of the standardization adopted in the Pearson correlation case. Indeed, in case standardization is implemented prior to the application of the two similarity indices, we get *J* = −0.426 and *C* = −0.320. Indeed, the similarity indices start behaving in similar qualitative manner to the Pearson when the colocalizations are preliminary standardized, which allows portions of the concentrations to become negatively interrelated (negative joint variations). These results again illustrate that quite distinct results can be obtained depending on the adopted normalization.

**Figure 21:**
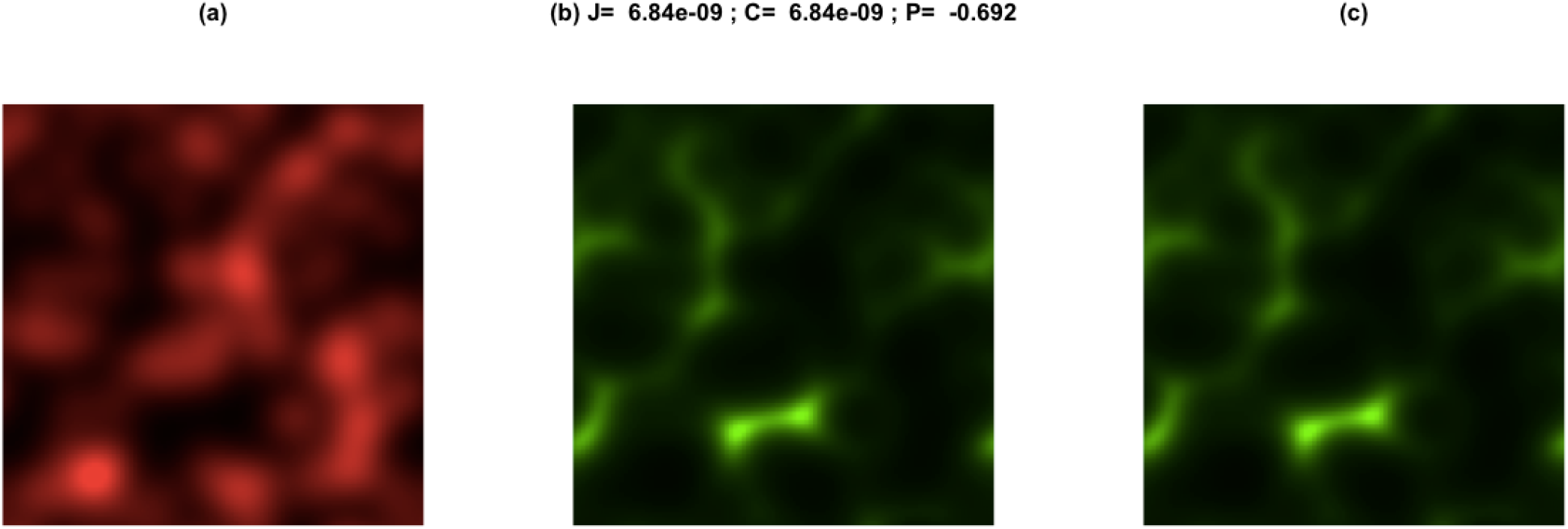
Example of colocalization analysis where one of the concentrations is the reciprocal of the other. Near zero results are observed for the similarity indices, with the Pearson correlation coefficient indicated negative relationship between the concentrations.

## 8 Concluding Remarks

Several real-world situations, especially in biology, involves signals that present similar spatial characteristics. The quantification of the colocalization between two or more concentrations corresponds to an important and frequently approach that has received substantial attention in the literature. Basically, the colocalization of two concentrations has been typically understood as relating to the tendency of the concentrations to vary together along space. By measuring a directly related tendency, the Pearson correlation coefficient has been frequently adopted for quantifying the colocalization in biological applications such as gene expression and protein synthesis.

Based on multiset theory, similarity indices have been described that are capable of quantifying the relationship between two generical mathematical structures, including concentrations, in terms of the relative shared area. The present work addressed the possibility to use the interiority, Jaccard and coincidence indices as a means to provide additional information about the colocalization of concentrations that can therefore complement the more traditional characterization in terms of the Pearson correlation coefficient.

We started by addressing the colocalization problem from a mathematical perspective of possible respective models, with emphasis on a model where a proportional iterrelationship is observe between the two concentrations. Several other modeling variations and possibilities were also discussed. These models can help not only better understanding the meaning of colocalization, but also help in choosing between normalizations as well as defining a more adequate manner to quantify the respective colocalization. The often critical issue of defining a baseline compatible with the poised questions was also characterized and discussed, given that this choice can have strong influence on the results obtained by the several considered methods of colocalization analysis.

One particularly important aspect that has been presented is the tendency of the Pearson correlation colocalization to be strongly biased by the presence of portions (peaks) of one of the concentrations that has no basic relationship whatsoever with the other concentration. In the considered respective example in this work, this effect resulted in a strong overestimation of the interrelationship (colocalization) between the two analyzed concentrations. This effect is mostly related to the displacement of the concentrations as implemented by the standardization. Indeed, the characterization of this type of concentrations by using the coincidence approach, which allows generic baselines (in this case that defined by the minmax normalization) to be taken into account. The zero baseline could then be considered, leading to a more meaningful colocalization quantification in this case.

We then presented the concepts of concentration average, which is essentially important respectively to the important concept of concentration baseline, as well as the standardization method, which is actually an intrinsic part of the Pearson correlation coefficient. Several examples of colocalization characterization by using this approach were the presented and discussed respectively to1D concentrations, which allow a more direct observation of the situations and effects.

The minmax normalization was then presented, followed by a description of the multiscale-based similarity indices.

It should be kept in mind that the choice of baseline, normalization, and indices for quantification of colocalization all can have strong influence on the obtained results. In addition, neither of them are absolute adequate or inadequate, as illustrated by several of the discussed examples. The choice between the several existing alternative should take into account how the data was generated, the specific ways in which the concentrations are intrinsically produced and interrelated by the systems of interest, as well as the questions implied by each respective research. A particularly interesting general approach is to consider several combinations of normalization and colocalization quantification approaches so that they can provide respectively complementary information about the prob-lem of interest, but even so the respective interpretation should take into account the above observed requirements and constraints.

By presenting new concepts and methods to be possibly applied for colocalization quantification, the present work paved the way to a large number of possible future developments. These include but are not limited to evaluating the proposed approaches respectively to more types of concentrations, considering alternative normalization methods, and extending the several described approaches to situations involving more than two concentrations.

## Acknowledgments

Luciano da F. Costa thanks CNPq (grant no. 307085/2018-0) and FAPESP (grant 15/22308-2).

## References

[1] Bolte S. and Cordelieres F. P. A guided tour into subcellular colocalization analysis in light microscopy. J. Microsc., 224:213–232, 2006.

[2] K. W. Dunn, M. M. Kamocka, and J. H. McDonald. A practical guide to evaluating colocalization in biological microscopy. Cell Physiology, 300(4):C723– C742, 2011.

[3] Zinchuk V. and Zinchuk O. A guided tour into subcellular colocalization analysis in light microscopy. Curr. Protoc. Cell Biol., 4(4:19), 2008.

[4] Oheim M. and Li D. Quantitative colocalisation imaging: concepts, measurements, and pitfalls. In Imaging Cellular And Molecular Biological Functions, page 449. Springer, Berlin, 2007.

[5] S. V. Costes, D. Daelemans, E. H. Cho, Z. Dobbin, G. Pavlakis, and S. Lockett. Automatic and quantitative measurement of protein-protein colocalization in live cells. Biophysical J., 6(6):3993–4003, 2004.

[6] J. Hein. Discrete Mathematics. Jones & Bartlett Pub., 2003.

[7] D. E. Knuth. The Art of Computing. Addison Wesley, 1998.

[8] W. D. Blizard. Multiset theory. Notre Dame Journal of Formal Logic, 30:36—66, 1989.

[9] W. D. Blizard. The development of multiset theory. Modern Logic, 4:319–352, 1991.

[10] P. M. Mahalakshmi and P. Thangavelu. Properties of multisets. International Journal of Innovative Technology and Exploring Engineering, 8:1–4, 2019.

[11] D. Singh, M. Ibrahim, T. Yohana, and J. N. Singh. Complementation in multiset theory. International Mathematical Forum, 38:1877–1884, 2011.

[12] M. K. Vijaymeena and K. Kavitha. A survey on similarity measures in text mining. Machine Learning and Applications, 3(1):19–28, 2016.

[13] P. Jaccard. Distribution de la flore alpine dans le bassin des dranses et dans quelques régions voisines. Bulletin de la Société vaudoise des Sciences Naturelles, 37:241–272, 1901.

[14] P. Jaccard. Étude comparative de la distribution florale dans une portion des alpes et des jura. Bulletin de la Société vaudoise des Sciences Naturelles, 37:547–549, 1901.

[15] B. K. Samanthula and W. Jiang. Secure multiset intersection cardinality and its application to jaccard coefficient. IEEE Transactions on Dependable and Secure Computing, 13(5):591–604, 1989.

[16] Wikipedia. Jaccard index. https://en.wikipedia.org/wiki/Jaccard_index. [Online; accessed 10-Oct-2021].

[17] A. Schubert and A. Telcs. A note on the Jaccardized Czekanowski similarity index. Scientometrics, 98:1397–1399, 2014.

[18] L. da F. Costa. Further generalizations of the Jaccard index. https://www.researchgate.net/publication/355381945_Further_Generalizations_of_the_Jaccard_Index, 2021. [Online; accessed 21-Aug-2021].

[19] L. da F. Costa. On similarity.https://www.sciencedirect.com/science/article/pii/S037843712200334X, 2022. Physica A: Statistical Mechanics and its Applications, 127456.

[20] L. da F. Costa. Multiset neurons. https://www.researchgate.net/publication/356042155_Multiset_Neurons, 2021.

[21] L. da F. Costa. Coincidence complex networks. https://iopscience.iop.org/article/10.1088/2632-072X/ac54c3, 2022. J. Phys.: Complexity, (3): 015012.

[22] L. da F. Costa. Comparing cross correlation-based similarities. https://www.researchgate.net/publication/355546016_Comparing_Cross_Correlation-Based_Similarities, 2021. [Online; accessed 21-Oct-2021].

[23] R. A. Johnson and D.W. Wichern. Applied multivariate analysis. Prentice Hall, 2002.

[24] F. Gewers, G. R. Ferreira, H. F. Arruda, F. N. Silva, C. H. Comin, D. R. Amancio, and L. da F. Costa Costa. Principal component analysis: A natural approach to data exploration. ACM Comput. Surv., 54(4):200–219, 2020.

[25] L. da F. Costa. Shape Classification and Analysis: Theory and Practice. CRC Press, Boca Raton, 2nd edition, 2009.

[26] D. P. Bertsekas and J. N. Tsitsiklis. Introduction to Probability. Athena Scientific, 2008.

[27] M. H. DeGroot and M. J. Schervish. Probability and Statistics. Pearson, 2011.

[28] E. Kreyszig. Advanced Engineering Mathematics. Wiley and Sons, 2015.

